# Nucleic Transformer: Deep Learning on Nucleic Acids with Self-attention and Convolutions

**DOI:** 10.1101/2021.01.28.428629

**Authors:** Shujun He, Baizhen Gao, Rushant Sabnis, Qing Sun

## Abstract

Much work has been done to apply machine learning and deep learning to genomics tasks, but these applications usually require extensive domain knowledge and the resulting models provide very limited interpretability. Here we present the Nucleic Transformer, a conceptually simple but effective and interpretable model architecture that excels in a variety of DNA/RNA tasks. The Nucleic Transformer processes nucleic acid sequences with self-attention and convolutions, two deep learning techniques that have proved dominant in the fields of computer vision and natural language processing. We demonstrate that the Nucleic Transformer can be trained in both supervised and unsupervised fashion without much domain knowledge to achieve high performance with limited amounts of data in *Escherichia coli* promoter classification, viral genome identification, and degradation properties of COVID-19 mRNA vaccine candidates. Additionally, we showcase extraction of promoter motifs from learned attention and how direct visualization of self-attention maps assists informed decision making using deep learning models.

---

DNA and RNA are essential components of life. They are associated with almost every biological process for eukaryotes and prokaryotes. Understanding DNA and RNA, including the stored information and properties such as degradation, is critical to understanding life and to reengineering biological systems [1]. However, DNA and RNA tasks have always been challenging because of the complex and large potential sequence space of DNA and RNA. Especially, the functions and properties of many coding and non-coding DNA/RNA sequences still remain poorly understood [2, 3].

Deep learning is a class of data-driven modeling approaches that have found much success in many fields including image recognition [4], natural language processing [5, 6], and computational biology [7]. It has allowed researchers to use recurrent neural networks (RNN/LSTM/GRU) to efficiently predict the function, origin, and properties of DNA/RNA sequences by training neural networks on large datasets [8, 9, 10, 11, 12, 13, 14, 15, 16, 17, 18], but these sequential computational approaches is difficult to parallelize and objects at long distances suffer from vanishing gradients. Transformers, on the other hand, is a recently proposed architecture that solely relies on attention mechanisms that can model dependencies regardless of the distance in the input or output sequences [5]. Although transformers have been adopted in many natural language processing tasks and seen massive success [5, 6, 19], we have found few studies using transformer in biological systems such as DNA/RNA sequences.

In this study, we present a model architecture Nucleic Transformer that utilizes convolution and self-attention to capture both local and global dependencies, which enable the model to achieve high accuracy and provide interpretability for three DNA/RNA tasks. The Nucleic Transformer formulates DNA understanding as natural language processing tasks, and RNA degradation as molecular prediction tasks. First, Nucleic Transformer is trained to classify short pieces of DNA sequence (81 bp) as either an *Escherichia coli* promoter sequence or non-promoter sequence. Nucleic Transformer learns from labeled promoter/non-promoter sequences, and it outperforms other state-of-the-art promoter identification models [20, 21, 22]. Second, Nucleic Transformer is tested on longer DNA sequences (300 bp) for classification of viral and non-viral sequences. We show that Nucleic Transformer predicts both viral and non-viral sequences with better accuracy compared with the previous best computational model [23]. In addition, we demonstrate that a Nucleic Transformer variant can be trained to predict RNA degradation rates at each position of a given sequence, a task of great importance to predict and produce stable mRNA vaccines and therapeutics.

## Results

### Nucleic Transformer combines self-attention and convolution to learn from DNA/RNA datasets

Any DNA seqeuence is a series of nucleotides, each of which can be one of A (adenosine), C (cytidine), G (guanosine), and T (thymine). Therefore, if we consider DNA a langauge that uses only four different characters to encode information, we can model it in similar fashion as a natural language. This idea then allows us to agnostically apply Natural Language Processing (NLP) techniques in the domain of deep learning without injection of domain specific knowledge in biology. Take the English language for example, all words in the vocabulary are combinations of the 26 letters in the alphabet; similarly, a DNA sequence is a sequence of 4 nucleotides. Although there are similarities between DNA sequences and natural language, there are also some important differences. The English langauge not only contains letters but also spaces that separate words and commas and periods that seperate sentences, whereas comparatively a DNA sequence is simply a sequence of 4 nucleotides. Further, when humans interpret a sentence, words are discretely recognized and each word can be considered a discrete object. As a result, state of art NLP models evaluate languages as collections of words (and punctuations) instead of letters. Since a DNA sequence does not have punctuations, we need to find a way to transform a DNA sequence into “words”. To do this, we transform the DNA sequences into k-mers with 1D convolutions (Figure 1a), a deep learning operation that allows for efficient and effective motif recognition.

**Figure 1:**
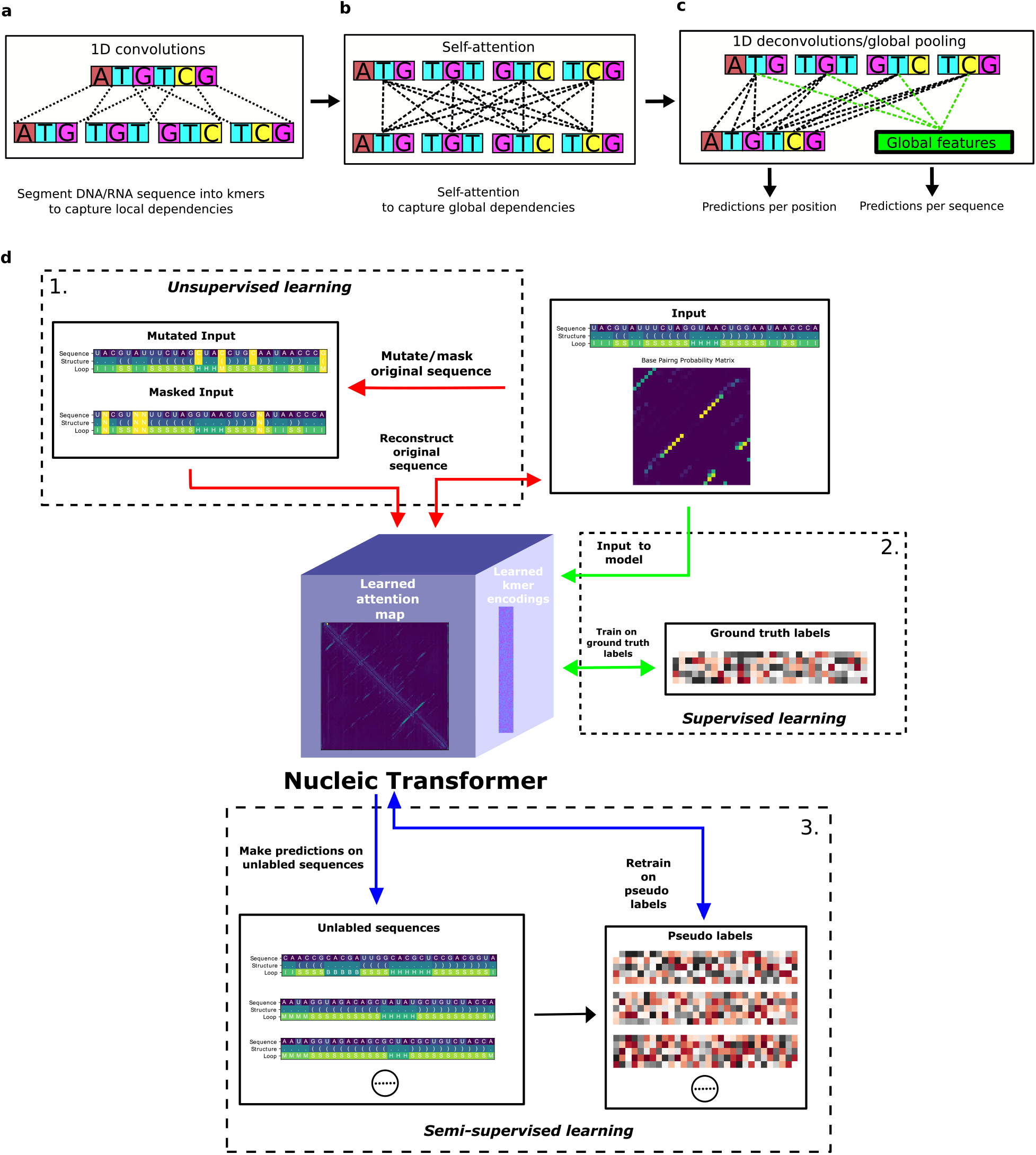
Nucleic Transformer combines self-attention and convolution to learn from DNA/RNA datasets. **a.** 1D convolutions is used on the nucleotide sequence to segment into k-mers **b.** Learn long-range dependencies are learned by self-attention **c.** Global pooling/deconvolution proceeds the transformer encoder for sequence level and nucleotide level predictions **d.** The Nucleic Transformer can be trained in unsupervised, supervised, and semi-supervise fashion.

Although effective at motif recognition, convolution on its own cannot capture long-range dependencies that are prevalent in DNA/RNA. Other works in the literature have applied recurrent neural networks such as LSTM (Long short term memory) networks, but LSTM’s sequential nature means that it is still insufficient at modeling relationships at long distances. To enable our neural network to learn long-range dependencies at any distance, we use transformer encoder layers with self-attention following the 1D convolution layers (Figure 1b). Self-attention networks, which process the entire sequence altogether, allow for faster computation compared to other sequence processing methods such as recurrent neural networks. Following the transformer encoder layers, we use one output layer (global pooling/deconvolution followed by a linear layer) right before making predictions appropriate for the task (Figure 1c). Combining 1D convolutions and self-attention makes our neural networks adept at modeling both local motifs and long-range dependencies, two key aspects of modeling DNA/RNA.

Additionally, we demonstrate that our neural network architecture can learn with different learning approaches, depending on the task at hand. Importantly,we highlight three key learning methods (Figure 1d):

1. Supervised learning, where the neural network learns from labeled input
2. Unsupervised learning, where the neural network learns directly from unlabeled input
3. Semi-supervised learning, where the training combines a small amount of labeled data and a larger amount of unlabeled data

### Nucleic Transformer accurately classifies DNA promoters and identifies consensus promoter motifs

We first demonstrate that the Nucleic Transformer outperforms previous results in the literature in *E. coli* promoter classification (Figure 2a). The Nucleic Transformer beats non-deep learning approaches which use sophisticated hand crafted features [24] by at least 1.7% or more. A more recent model, iPromoter-BnCNN [25], which also employs structural property properties of DNA such as stability, rigidity, and curvature, comes close in performance to the Nucleic Transformer, although the Nucleic Transformer directly makes predictions from sequence information. Further, to compare transformer encoder with LSTM, we swapped out the transformer encoder with a Bidirectional LSTM while keeping all other conditions the same. We see that using transformer encoder consistently outperforms LSTM on all *k* values (Figure S1a). On the independent test set (Figure 2b), which includes recently released experimentally verified promoter samples, the Nucleic Transformer is more accurate than MULTiPly, iPromoter-2L, and iPromoter-BnCNN [21, 24, 25].

**Figure 2:**
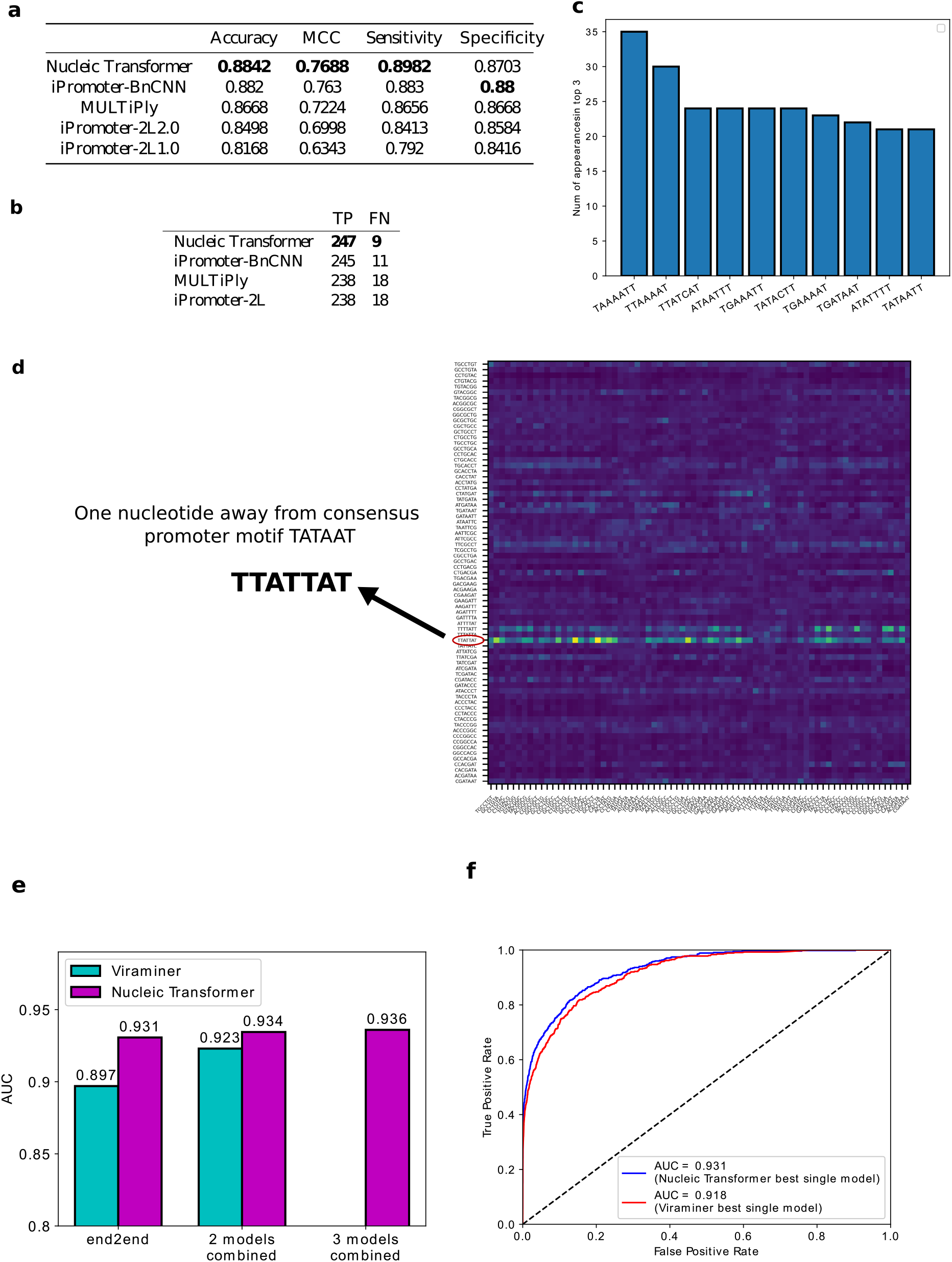
Nucleic Transformer outperforms previous top results in DNA classification. **a.** Performance of Nucleic Transformer against top results in the literature based on accuracy, MCC, sensitivity, and specificity. **b.** Performance of Nucleic Transformer on the independent test set against top results in the literature. **c.** Most important 7-mers in promoter classification based on analysis of attention weights. **d.** Visualization of an attention weight right before outputting a prediction. A bright band can be seen for the 7-mer TTATTAT, which is one mutation away from TATAAT. **e.** Comparison of test set performance between Nucleic Transformer and Viraminer. **e.** AUC plot comparing test set performance between best single end to end models of Nucleic Transformer and Viraminer.

By visualizing the attention weight matrix, we see that the Nucleic Transformer often focuses on kmers that closely resemble consensus promoter motifs (Figure 2d). We also extracted motifs that the Nucleic Transformer considers the most characteristic of promoters (Figure 2c). For each correct prediction of promoters in the validation sets of 5-fold cross validation, we take the column-wise sum of the attention weight matrix and rank the kmers. Then we simply count the amount of times each kmer receives top 3 most attention in all correct predictions. We found the kmers that frequently appear in top 3 resemble the consensus promoter motif TATAAT. In fact, one of the 10 most frequently 7-mers have the exact consensus motif TATAAT, while 6 others contain the motif TAAAAT, TATCAT, ATAAT, TATACT, GATAAT, all of which are one mutation away from the consensus motif TATAAT. Deep learning has been criticized as black boxes that cannot be interpreted [26]; however, here we demonstrate that the Nucleic Transformer can be interpreted and useful motifs can even be derived from attention weights directly.

### Nucleic Transformer outperforms previous models in viral genome classification

To further demonstrate the effectiveness of the Nucleic Transformer architecture, we trained Nucleic Transformer models on a viral/non-viral genome dataset previously used to train Viraminer, a purely convolution based model [23], and compare the performance of the two models. When trained end to end, the Nucleic Transformer significantly outperforms the Viraminer counterpart by 3.4% (Area Under the ROC Curve) AUC score (Figure 2e). When trained with a two-stage training process combining two branches with different hyperparamters and pooling schemes, the Viraminer performs significantly better compared to its end to end counterpart, but the Nucleic Transformer still leads in accuracy even with just one end to end model (*k* = 13). It is widely known that even just simply averaging results from a few machine learning models trained with slightly different hyperparameters usually leads to better performance, so it is no surprise that Viraminer’s strategy to combine two branches with different setups proves successful. For better comparison, we also trained the Nucleic Transformer 2 and 3 times with different *k*’s and averaged the test set predictions. The AUC improved upon by 0.3% upon averaging 2 models (*k* = 11, 13), but with 3 models averaged (*k* = 11, 13, 15), the AUC only improved slightly by 0.2%, which is no no surprise since with more models ensembled there is usually diminishing returns. We also compare the ROC curve between the best single models of Viraminer (Frequency branch) and the Nucleic Transformer (Figure 2f) and we see that the Nucleic Transformer holds a clear advantage in single model performance.

### Nucleic Transformer accurately predicts RNA degradation at nucleotide level

mRNA vaccines and therepeutics have incredible potentials, but one key limitation is the inherent instability of mRNA. The 21-day OpenVaccine challenge, hosted by Das Lab at Stanford University [27], sought to understand RNA degradation by rallying the expertise of data scientists from all over the globe. We used an adaption of the Nucleic Transformer in this challenge with some key modifications. Unlike DNA which is double stranded and relatively stable, RNA is single-stranded and highly promiscuous. While double-stranded DNA forms hydrogen bonds between its complementary bases, single-stranded RNA forms secondary structures by itself, which have been known to stablize RNA molecules. Therefore, we use existing biophysical models to inject knowledge of RNA secondary structure into our deep learning models to predict RNA degradation. To do so, we simply add the base pairing probability map following 2D convolutional layers as an additional bias to the self-attention weight in self-attention layers (Figure 3a), while adding additional embedding layers to represent structure and loop predictions from secondary structure models.

**Figure 3:**
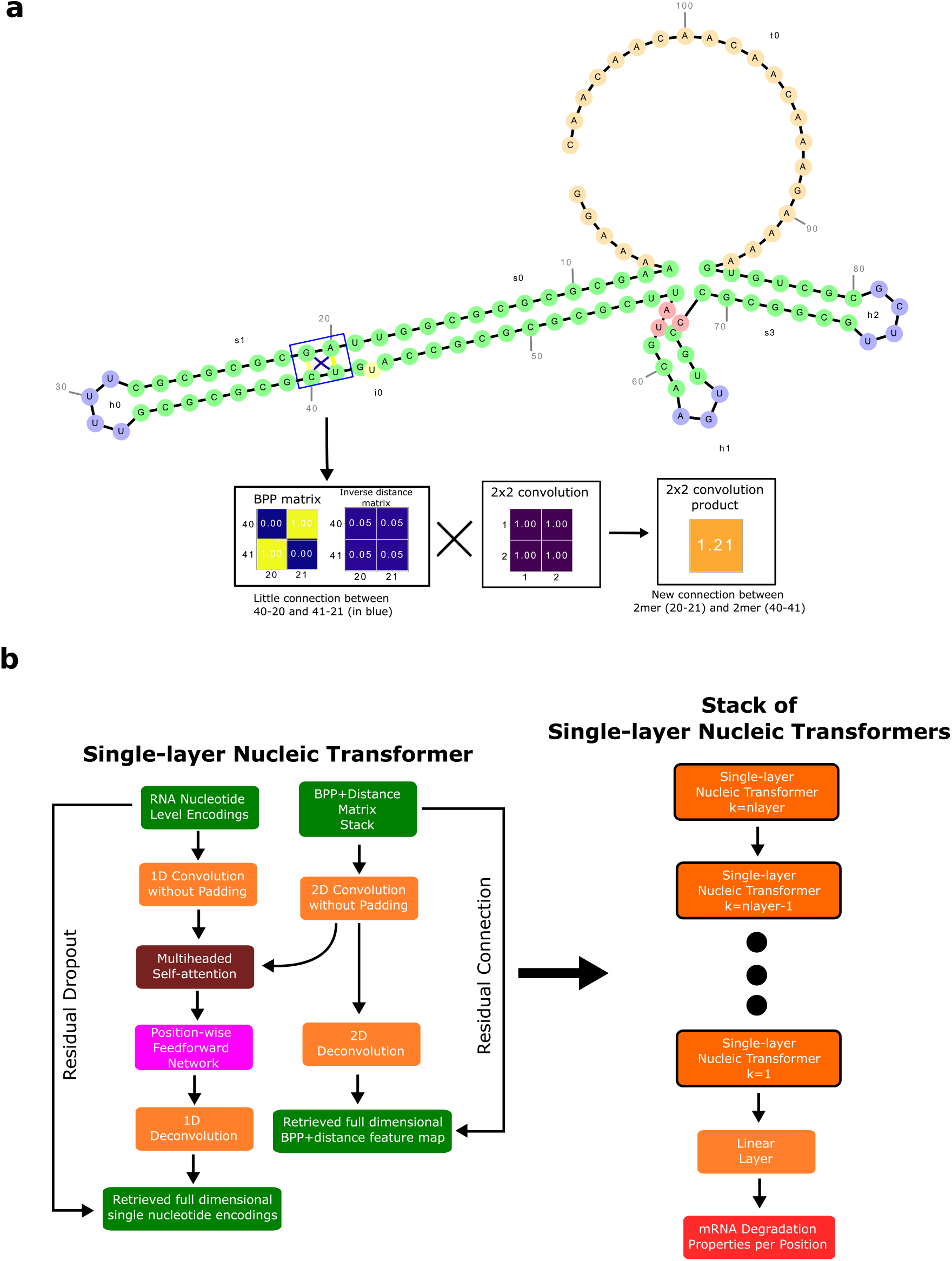
Nucleic Transformer for RNA degradation. **a.** Visualization of BPP+distance matrix, attention weights of a non-pretrained Nucleic Transformer, and attention weights of a pretrained Nucleic Transformer **b.** Nucleic Transformer stack which takes advantage of additional input information from biophysical models.

#### Interpretiblility of self-attention assists decision making

The test dataset in OpenVaccine was split into a public and private set. The score on the public test set was available during the competition, while the score on the private test set was only revealed at the end of the competition and used for the final ranking. Notably, the public set is composed of 107 bp RNA sequence stability data while private set consists of more diverse and longer (137 bp) mRNA sequences. Each team was afforded two submissions to be used for the final ranking. In machine learning competitions, competitors would normally have to rely on local cross validation to evaluate model performance. Nevertheless, the training set in OpenVaccine had noisy training samples and limited sequence diversity, which meant that even local cross validation may not be robust enough. Therefore, we had to look for other avenues to robustly select the best models for the final ranking.

By the end of the competition, we had trained two sets of models to use in the final submissions, one that was trained directly on short sequences with labels and one that was pretrained with all available sequences (including unlabeled test sequences) before training on short sequences with labels. In order to robustly select submissions, we visualized and evaluated the learned attention weights from the transformer encoder (Figure 4a). Since we added the base pairing probability (BPP) matrix and distance matrix as a bias, both learned attention distributions of pretrained and non-pretrained models resembled the BPP and distance matrix, but there were also some key differences. The non-pretrained model paid heavy attention indiscriminately to pairs of positionally close nucleotides, as indicated by the bright stripes parallel and close to the diagonal of the attention matrix. This indicates the non-pretrained model thought that positionally close nucleotides were always important when making predictions on mRNA degradation properties, which seemed highly unlikely. On the other hand, the pretrained model did not show the same bias towards pairs of positionally close nucleotides, and was able to recognize the weak BPP connections which were barely visible on the original BPP matrix. In this case, the model made more effective use of the BPP matrix generated by biophysical models. Because of these considerations, we favored pretrained models in our final submissions, where one submission was an average of 20 pretrained models, and the other was an average of 20 pretrained models and 20 non-pretrained models.

**Figure 4:**
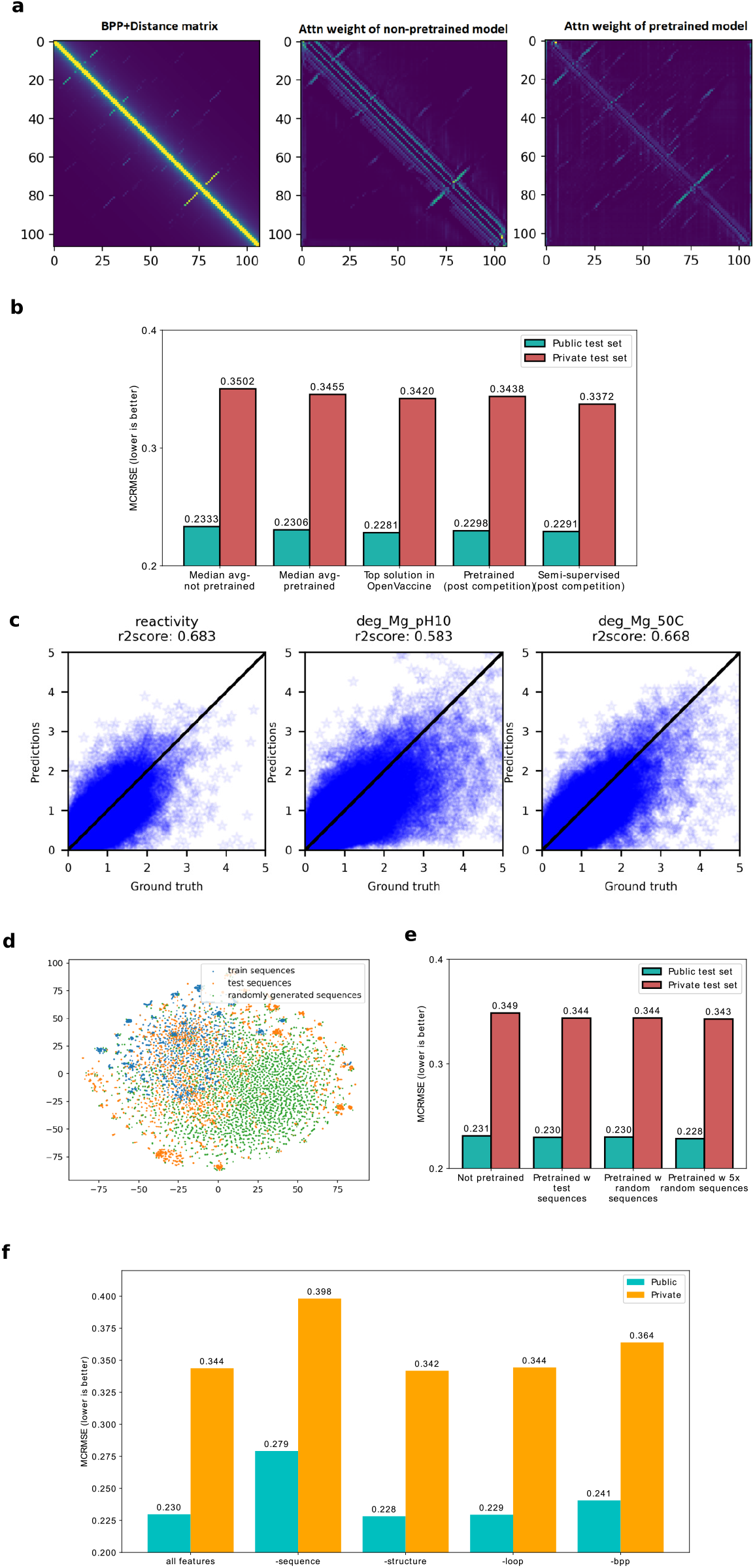
Nucleic Transformer accurately predicts RNA degradation at nucleotide level. **a.** Visualization of BPP+distance matrix, attention weights of a non-pretrained Nucleic Transformer, and attention weights of a pretrained Nucleic Transformer **b.** Comparison of pretrained and non-pretrained models trained during the OpenVaccine competition and post competition experiments. **c.** R2 plot of best ensemble of semi-supervised learning results on the private test set. **d.** Tsne plot of RNA sequences in the training set, test set, and randomly generated set. **e.** Test set performance of OpenVaccine challenge using different pretraining procedures. **f.** Leave one feature out (LOFO) feature importance on the OpenVaccine dataset.

Results on the private test set validated our selection based on visual inspection of attention weights (Figure 4b). Pretrained models performed significantly better than non-pretrained models on both public test set and private test set. The non-pretrained models would have placed us at 39th/1636 instead of 7th/1636. Notably, the predictions on the private test set have much higher error, likely both due to longer sequence length and more sequence diversity in the private test set.

#### Semi-supervised learning leads to more accurate predictions in RNA degradation

Following the competition, we found that using predictions generated by an ensemble of 5 Nucleic Transformers as pseudo-labels, we could reduce the test set MCRMSE to 0.33722, compared to the top solution’s 0.34198 (Figure 4b). This is somewhat surprising given the that the predictions used as pseudo-labels could only score 0.3438 on the private test set. When we compare the R2 scores of supervised only, unsupervised, and semi-supervised learning, we see that unsupervised and semi-supervised resulted in significant improvements over the supervised only (Figure 4c). Also, we notice that predictions for deg_Mg_pH10 have significantly larger errors than the other 2 properties, which is expected because the biophysical models used are incapable of generating different secondary structure predictions at different pHs. It is also important to note that the semi-supervised learning approach requires pseudo-labeling and knowing the test set distribution before hand. Therefore, while it is effective under the competition setting, it is a risky approach and the performance gain may not transfer in real life applications.

#### Learning from randomly generated sequences

To explore whether the pretraining performance gain results from knowing the test set distribution, we also generated additional completely random sequences to use for pretraining. While the sequences in the test set have more sequence diversity than the training set, randomly generated sequences are even more diverse (Figure 4d). We repeated the pretraining process with random sequences using the same hyperparamters, and our experiments show that pretraining with random sequences instead of test sequences results in almost identical performance and increasing the amount of random sequences for pretraining leads to slightly better performance on both test sets (Figure 4e). These results suggest that pretraining improves test set error not because of information leak from knowing the test sequences but rather the model learning the generalized rules of mRNA secondary structure formation. Additionally, we also pseudo-labeled random sequences and retrained our models in semi-supervised fashion. In this case, pseudo-labeling with completely random sequences did not lead to significant improvements. It is likely that due to the large mutation distance of random sequences compared to train and test sequences, the pseudo-labels on the random sequences do not provide useful information. Therefore, the semi-supervised learning process ended up feeding mostly noise to the model during training, although it did lead to very slight improvement on the private test set over the model without semi-supervised learning.

#### Most important features in RNA stability predictions

To understand what features are the most important in predicting RNA stability, we took a simple LOFO (leave one feature out) approach. We retrained our models from scratch with the best settings but leave out one of the available features (sequence, predicted structure, predicted loop type, or predicted base pair probabilities), and then evaluated model performance with the absence of one feature (Figure 4f). We found that our models took the biggest performance hit when sequence information was left out, followed by predicted base pair probabilities. Surprisingly, leaving out structure and loop features had little impact on model performance as the MCRMSE remained almost the same. Our hypothesis is that the structure and loop features represent the most likely fold based on predicted base pair probabilities, while in reality, mRNA is supposed to have an ensemble of folds, so in this case the base pairing probability matrix provides richer and more detailed information and is more important.

## Discussion

In this work, we present the Nucleic Transformer, an effective and yet conceptually simple architecture that is capable of high performance in classifying promoters and viral genome as well as predicting degradation properties of mRNA COVID-19 vaccine candidates per nucleotide. The Nucleic Transformer architecture outperforms other deep learning/non deep learning methods that require hand-crafted features in *E. coli* promoter classification, while also providing interpretibility and being capable of extracting promoter motifs directly from learned attention. Although always trained end to end, the Nucleic Transformer has better accuracy in classifying viral genomes compared to previous models such as Viraminer, which requires sophisticated multi-stage training and ensembling. As an additional test, we also participated in the recent OpenVaccine challenge with an adaptation of the Nucleic Transformer and placed 7th out of 1636 teams of top machine learning experts from all over the globe. We also demonstrate with semi-supervised learning, the Nucleic Transformer outperforms even the top solution in the OpenVaccine challenge by a considerable margin. Our results show that self-attention and convolution are a powerful combination for genomics tasks, enabling learning both global and local dependencies effectively. It has long been posited that the transformer architecture can excel beyond natural language processing, and our work demonstrates that.

Although classification of promoters and viral genomes have been well studied in the literature, there is no precedence on predictions of mRNA degradation properties per nucleotide. After a chaotic 2020 caused by COVID-19, mRNA vaccines have emerged as a fast and effective solution to the COVID problem, with companies like Pfizer and Moderna rolling out mRNA vaccines at unprecedented speeds. However, storage and transport remain a challenge with fragile mRNA vaccines (Pfizer-BioNtech’s vaccine has to be stored as −80 °C and Moderna’s at −20 °C). One strategy to reduce mRNA hydrolysis is to redesign RNAs to code for the same proteins but form double-stranded regions, which are protected from these degradative processes [28]. Our work can provide guidance and trained models can act as a screening tool in development of more stable mRNA vaccines in the future. mRNA vaccines and therapeutics have many applications towards infectious diseases and cancers [29, 30, 31, 32], and it is our hope the Nucleic Transformer will aid design of more stable mRNA vaccines that can withstand harsher conditions than current ones. It is important to note, however, that there is still significant gap between errors on the 107 bp mRNA OpenVaccine public set sequences and the 130 bp mRNA OpenVaccine private set sequences, both due to difference in sequence length and diversity. Actual COVID-19 candidates are even longer and modeling those remains a challenge in the future. For these long sequences, one key challenge is the quadratic computational complexity of self-attention, which prohibits training on long sequences. Notably, much work has been done in very recent times on reducing the quadratic computational complexity of self-attention to linear to enable training of transformer like self-attention on much longer sequences [33, 34, 35, 36], i.e. linear transformers. Whole genomes and COVID-19 mRNA vaccines both greatly exceed the length limit of full self-attention, and COVID-19 vaccine candidates, in particular, are around 4000 bp long, so these new approaches may serve as effective tools to solve the challenges.

In summary, we have developed a convolution and transformer based deep learning platform towards new and urgent DNA/RNA tasks. Our work has demonstrated success in DNA sequence identification and RNA stability predictions. We believe that by further development and optimization, we will solve many nucleotides challenges including understanding RNA decay and structure relationships, aiding next generation mRNA therapy development, reading out information from unknown DNA and RNA sequences, and more.

## Methods

### Benchmark datasets

#### *E. coli* promoter/non-promoter dataset

The *E. coli* promoter/non-promoter dataset is an experimentally confirmed benchmark dataset widely used in the literature to model and evaluate DNA promoter sequences [24]. All DNA sequences in the dataset were collected from RegulonDB, and sequences were screened by CD-HIT based on redundant sequence identity [37]. This dataset consists of 2,860 promoter sequences and 2,860 non-promoter sequences. All promoter sequences were experimentally confirmed and collected from RegulonDB (version 9.3) [38]. The non-promoters sequences were extracted randomly from the middle regions of long coding sequences and convergent intergenic regions in *E. coli* K-12 genome [39, 40]. Model performance on this dataset is evaluated using 5-fold cross validation, where the data is split using iterativte stratification [41]. The metrics used for this dataset are accuracy, sensitivity, specificity, and Matthews correlation coefficient (MCC). In addition to cross-validation, we also use an independent test set composed of experimentally verified *E. coli* promoters recently added to RegulonDB.

#### Viral/non-viral genome dataset

The viral/non-viral genome dataset is same as the one used to trained viraminer [23], which consists of 19 different NGS experiments analyzed and labeled by PCJ-BLAST [42] following de novo genome assembly algorithms. This dataset is publicly available at https://github.com/NeuroCSUT/ViraMiner. DNA sequences included in this dataset were cut to 300 bp segments with remaining portions smaller than 300 bp discarded. Further, all sequences that contain “N” (unknown with equal probability to any of the four nucleotides) were removed as well. This dataset has approximately 320,000 DNA sequences in total. The main challenge with this dataset is the class imbalance, where only 2% of sequences are of viral origin. The dataset is split into training, validation, and test, where hypertuning was done with the training and validation set, and model performance evaluated on the test set. The metric used for this dataset is AUC (Area Under the Receiver Operating Characteristic Curve).

#### OpenVaccine challenge dataset

The OpenVaccine challenge [27] hosted by the Eterna community sought to rally the data science expertise of Kaggle competitors to develop models that could accurately predict degradation of mRNA. During the 21-day challenge, competitors were provided with 2400 107-bp mRNA sequences with the first 68 base pairs labels with 5 degradation properties at each position. These properties are reactivity, deg_pH10, deg_Mg_pH10, deg_50C, and deg_Mg_50C. More details on these properties can be found at https://www.kaggle.com/c/stanford-covid-vaccine/data.

Like most Kaggle competitions, the test set was divided into a public test set and a private test set. Results on the public test set was available during the competition, while private test set results were hidden. The final evaluation was done on a portion of the private test set consisting of 3005 130-bp mRNA sequences, whose degradation measurements were conducted during the 21-day challenge and revealed at the end. The test set was subjected to screening based on three criteria:

1. Minimum value across all 5 degradation properties must be greater than −0.5
2. Mean signal/noise across all 5 degradation properties must be greater than 1.0. [Signal/noise is defined as mean(measurement value over 68 nts)/mean(statistical error in measurement value over 68 nts)]
3. Sequences were clustered into clusters with less than 50% sequence similarity and chosen from clusters with 3 or fewer members

After screening, only 1172 sequences remained on the test set. Final evaluation was done on 3 of the 5 properties (reactivity, deg_Mg_pH10, and deg_Mg_50C). Unlike the training set, the test set has longer mRNA sequences with more sequence diversity and more measurements (first 91 positions) per sequence; in fact, more predictions had to be made for the test set than there were training samples. The OpenVaccine challenge is thus the most challenging among the tasks tackled in this paper. The metric used for ranking in the competition is MCRMSE (mean columnwise root mean squared error):

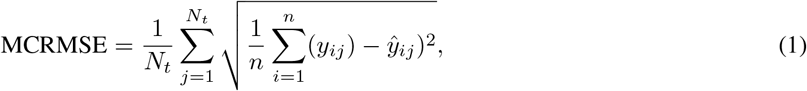

where *N_t_* is the number of columns, *n* the number of positions predicted, *y* the ground truth, and *y* the predicted value. In addition, we also use R2 score (coefficient of determination) during further analysis.

### Architecture design

#### K-mers with 1-D convolutions

The extraction of k-mers from a DNA sequence can be considered a sliding window of size *k* taking snapshots of the sequence while moving one position at a time from one end of the sequence to the other, which is conceptually identical to the convolution operation used in deep learning. Consider a simple example of convolution involving a vector 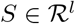, where *l* is the length of the vector, and a convolution kernel 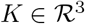, which convolves over the vector *S*. If the convolutional kernel strides one position at a time, an output vector of dot products 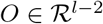 is computed,

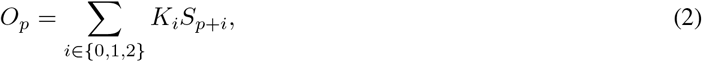

where *p* denotes the position in the output vector. In this case, the convolution operation aggregates local information with 3 positions at a time, so if *S* is a sequence of nucleotides, the convolution operation is essentially extracting 3-mers from the DNA sequence *S*.

Since our model takes sequences of DNA/RNA nucleotides as input, we first transform each nucleotide into embeddings of fixed size *d_model_*. So now for each sequence we have a tensor 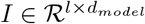, where *l* is the length of the sequence. Because we are using the transformer encoder architecture, which is permutation invariant, we need to add positional encoding, same as the implementation in [5]:

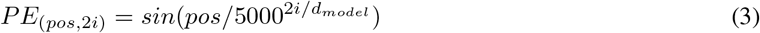

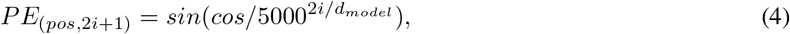

where *pos* is the position and *i* is the channel dimension. Now to create k-mers we perform convolutions on the tensor *I* without padding and stride = 1. When a convolution operation with kernel size *k* is performed over *I*, a new tensor 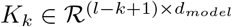 representing the sequence of k-mers is generated. Now each k-mer is represented by a feature vector of size *d_model_*. The 1D convolution layers are always followed by a layer normalization layer [43].

Indeed our representation of k-mers deviates from conventional representation of words in deep learning, where each word in the vocabulary directly corresponds to a feature vector in a look up table. The disadvantage of using look up tables for k-mers is that a very small percentage of all possible k-mers are present in any dataset, and there is no way for the network to generalize to unseen k-mers. For instance, if a common promoter motif TATAAT appears in the dataset but a similar motif TATATT does not, there is no way for the network to generalize to the unseen motif TATATT, but by using convolutions, we make it easy for the network to recognize that there is not much difference between TATAAT and TATATT, leading to better generalization. Additionally, embeddings of k-mers of larger sizes require a prohibitively large amount of parameters, since the total possible amount of k-mers for a given *k* is 4^*k*^.

#### Transformer encoder

For the encoder, we implement the vanilla transformer encoder [5], which uses the multi-head self-attention mechanism. First, the k-mer representations (each kmer is represented by a feature vector of size *d_model_*) are linearly projected into lower dimensional (*d_model_*/*n_nhead_*) keys, values and queries for *n_nhead_* times. Next, the self-attention function is computed with the lower dimensional keys, values and queries for *n_nhead_* times independently. In this case, the self-attention function essentially computes a pairwise interaction matrix relating every k-mer to every k-mer (including self to self interaction) and computes a weighted sum of the values. It has been posited that the multi-head mechanism allows different heads to learn different hidden representations of the input, leading to better performance. The multi-head self-attention mechanism can be summarized in a few equations:

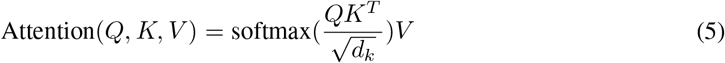

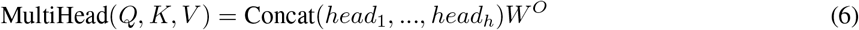

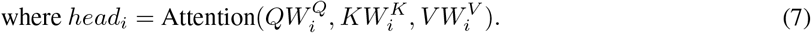

Since we are only using the transformer encoder, *Q*, *K*, *V* come from the same sequence of feature vectors (hence the name self-attention), each of which represents a k-mer with positional encoding.

The self-attention mechanism enables each k-mer to attend to all k-mers (including itself), so global dependencies can be drawn between k-mers at any distance. Contrary to recurrence and convolutions, both of which enforce sparse local connectivity, transformers allow for dense or complete global connectivity. The ability to draw global dependencies of transformers is a huge advantage over recurrence and convolutions, both of which struggle with long sequences.

The self-attention function is followed by a position-wise feedforward network applied separately and identically to each position:

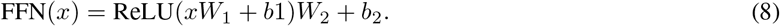

The position-wise feedforward network is basically two linear transforms with a ReLU activation in between. Conventionally, the combination of self-attention and position-wise feedforward network is referred to as the transformer encoder layer, and a stack of transformer encoder layers is referred to the transformer encoder.

#### Incorporating biophysical models to predict RNA degradation

Although we found it sufficient to simply use sequence information for DNA tasks, predicting degradation of RNA requires more than just sequence information. Here we detail the changes made to utilize biophysical models. Firstly, we include the predicted structure per position, which describes whether a nucleotide is paired or unpaired with another one via hydrogen bonding. The predicted structure is generated using arnie with log_gamma set to 0. Also, we include the predicted loop type assigned by bpRNA [44]. Two embedding layers are added to represent structure and loop type, and the resulting feature vectors are concatenated and the dimensionality reduced with a linear transformation. Additionally, we directly add a modified version of the base-pairing probability matrix into the attention function (note that this is simply a modified version of Eq.5 with *M_bpp_* as an additional input):

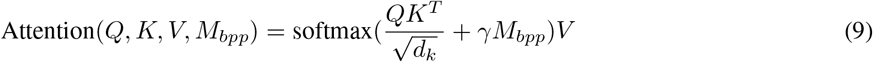

where *γ* is a learnable parameter and *M_bpp_* is the modified base-pairing probability matrix. The original base-pairing probability matrix contains the probabilities for every possible base pair in an RNA sequence and has been used for many RNA informatics tasks. Here in addition to base-pairing probabilities, we also stack inverse, inverse squared, and inverse cubed pairwise distance matrices on top of the original base-pairing probability matrix, where the distance is the the number of covalent bonds between the pair of nucleotides (this can also be considered the path length in an RNA graph where the only edges are the covalent bonds). The inverse distance matrices encode some information about the relative distance between pairs of nucleotides, since pairs of nucleotides with a small number of covelent bonds in between are likely to be closer to each other spatially. Because the distance matrix already encodes information about position, we do not use positional encoding for mRNA.

Because 1-D convolution operation used in the Nucleic Transformer does not use padding, the convolution product ends up with reduced dimensionality in the *L* dimension when the convolution kernel size is bigger than 1. As a result, the base pairing probability matrix cannot be directly added to self-attention matrix. To circumvent this, we also do 2D convolution with the same kernel size as the 1D convolution on the modified base pairing probability matrix without padding, so the dimensionality of the feature map becomes *C* × (*L* – *k* + 1) × (*L* – *k* + 1). The attention function now is:

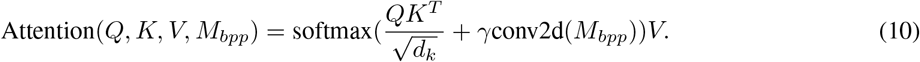

Conceptually, instead of a base pair to base pair interaction mapping, the 2D convolution product of the modified base pairing probability matrix can be seen as a kmer to kmer pairwise interaction mapping with matching dimensionality to the 1D convolution kmer products. Aside from matching dimensionality, the 2D convolution operation also makes up for some missing information regarding the geometry of mRNA folding. To illustrate this, we visualize an mRNA sequence in the OpenVaccine dataset to explain the physical and mathematical reasoning behind the 2D convolution operation (Figure 3b). While inspecting the interaction between A-20 (A at position 20), G-21, C-40, and U-41, we can visually see that A-20 and C-40 are quite close to each other and imagine that there is some degree of interaction between them, despite A-20 and C-40 not forming hydrogen bonds. However, looking at the portion of BPP matrix and distance matrix corresponding to the 2×2 connection between A-20 (A at position 20), G-21, C-40, and U-41, we see that neither the BPP matrix nor the distance matrix convey this information, as the component (40,20) has zero or close to zero values on both the BPP matrix and the distance matrix. When a 2×2 convolution kernel operates on the BPP matrix and distance matrix (for illustration purposes here we simply draw a kernel with all values set to unity), it essentially fuses the 4 connections between A-20, G-21, C-40, and U-41, and creates a strong connection between the 2 2mers (A-20, G-21 and C-40, U-41). Now it becomes much easier for the network to learn the interaction between A-20 and G-40 (as well as for G-21 and U-41).

The combination of convolution and self-attention cannot produce nucleotide position wise predictions, since it generates kmer encodings instead of single base pair encoding. In order to make predictions per nucleotide position, we introduce additional deconvolution layers to retrieve full dimensional encodings, which allow residual connections of both 1D and 2D encodings before and after the transformer encoder. As a result, both the single nucleotide embeddings and the modified BPP matrix go through deep transforms before outputting predictions.

Now we can summarize the modified Nucleic Transformer architecture used for the RNA task (Figure 3b), which can be seen as a special case of a series of multiple Nucleic Transformers with a single transformer encoder layer followed by a deconvolution layer. Also, because the OpenVaccine challenge requires making predictions at each position of the RNA sequence, it is important for the last transformer encoder layer right before outputting predictions to operate on single nucleotide encodings instead of kmer encodings. With these considerations in mind, we choose a simple strategy to construct the stack of Nucleic Transformers with two main hyperparameters *k* and *n_layer_* set equal to each other. The first single layer Nucleic Transformer has *k* = *n_layer_* and we decrease the size of the convolution kernel by 1 for the next single layer Nucleic Transformer. Therefore, when we get to the last Nucleic Transformer in the stack, *k* becomes 1 and the last Nucleic Transformer is simply a transformer encoder layer with an added bias from the BPP feature map.

### Training details

#### Optimizer and training schedule

For DNA classification tasks, we choose Adam [45], a commonly used optimizer in deep learning with *β*_1_ = 0.9, *β*_2_ = 0.99, and *ε* = 1e – 8. Weight decay is set to 1e-5. Our learning rate schedule is a stepwise inverse square root decay schedule with warm up. Since we use relatively small batch sizes during training, we adjust the learning rate by a scaling factor *C*:

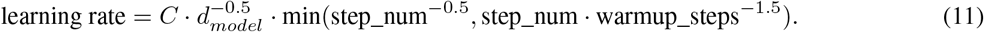

In our experiments, *C* is set 0.1 and warmup_steps is set to 3200. Additionally, we use dropout [46] of probability 0.1 in all attention layers, fully connected layers, and positional encoding.

For the RNA task, we found Adam to be underfitting. Therefore, we switched to a more recent and powerful optimizer, Ranger, which uses gradient centralization from https://github.com/lessw2020/Ranger-Deep-Learning-Optimizer [47]. As for the training schedule, we used flat and anneal, where training starts with a flat learning rate of 1e-3 and then 75% through all the epochs training proceeds with cosine annealing schedule reducing learning rate down to 0 at the end of training. Weight decay is set to 0.1.

#### Learning approaches

For DNA tasks, our models learn in supervised fashion. However, in the OpenVaccine challenge, the number of samples in the test set exceeds that in the training set, so we use a combination of learning methods (Figure 1d). Here we describe those learning methods.

##### Multitasking learning

during pretraining, mutated/masked sequence, structure, and predicted loop type are inputted into the Nucleic Transformer and then the Nucleic Transformer is trained with crossentropy loss to retrieve the correct sequence, structure, and predicted loop type simultaneously at each position of the RNA sequence. During training on ground truth labels and pseudo labels, the Nucleic Transformer is trained to predict 5 different degradation properties simultaneously at each measured position of the RNA sequence.

##### Supervised learning

during supervised learning, the Nucleic Transformer is trained on target values of classification classes or RNA degradation properties.

##### Unsupervised learning

we use all available sequences in the OpenVaccine challenge dataset to pretrain our network on randomly mutated and masked (with NULL token) sequence retrieval loss (basically softmax to retrieve correct nucleotide/structure/loop). During pretraining, the Nucleic Transformer learns the rules of RNA structure, guided by biophysical knowledge provided by biophysical models.

##### Semi-supervised learning

Following RNA supervised learning, the Nucleic Transformer is retrained in semi-supervised fashion on pseudo labels generated by an ensemble of Nucleic Transformers with different depths. Similar to previous work with semi-supervised learning [48], we retrain the models first using pseudo labels at a flat learning rate and then finetune with ground truth labels in the training set with cosine anneal schedule.

#### Error weighted loss

Because the OpenVaccine dataset came from experimental measurements that had errors, we adjusted the losses based on the error for each measurement during supervised training:

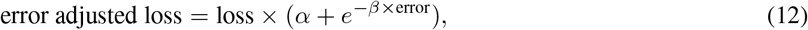

where *α* and *β* are tunable hyperparameters. If *α* is set to 1 and *β* to infinity, the loss values stay the same; otherwise gradients from measurements with large errors would be lowered to prevent the neural network from overfitting to experimental errors. For the OpenVaccine dataset, we use *α* = 0.5 and *β* = 5.

#### Usage of biophysical models during training

We used secondary structures predicted by an ensemble of biophysical models including RNAsoft [49], rnastructure [50], CONTRAfold [51], EternaFold [52], NUPACK [53], and Vienna [54]. Arnie^1^ is used as a wrapper to generate secondary structure predictions. For each sequence, we generated secondary structure predictions at 37 and 50 Celsius, since two of the scored degradation properties were measured at different temperatures. Although we also need to make predictions for a degradation property at pH10, none of the biophysical models used could generate predictions at different pH’s. With 6 packages, we ended with up with 12 secondary structure predictions for each sequence. During training, we randomly select one of the 12 secondary structure predictions for each sample during a forward and backward propagation pass. During validation and testing, we use the averaged predictions made with all 12 secondary structure predictions.

#### Random mutations during training

It is known that deep learning models can perfectly memorize completely random data and labels, so to combat the memorization effect, we inject noise artificially by randomly mutating positions in the source DNA/RNA sequence before forward and backward propagation during training, similar to bert’s pretraining [6]. This injection of random noise is done during DNA supervised learning and RNA unsupervised learning. Note that in all our experiments, we simply randomly mutate nucleotides in randomly selected positions, and because we do not ensure nucleotide at each selected position is changed, the average amount of mutations is 3/4 of *n_mute_*. It is true that these random mutations could be non-label-preserving; however, deep learning algorithm is robust to massive label noise, so the non-label-preserving mutations should simply be ignored by the network during training [55]. The number of positions to randomly mutate is a hyperparamter that we denote *n_mute_*, with which we experiment to find the best value.

### Best hyperparameters for different tasks

For promoter classification, the best results were obtained using *k* = 7, *n_mute_* = 15, six transformer encoder layers, *d_model_* = 256, *n_head_* = 8, and batch size is 24. For viral/non-viral DNA classification, the best results were obtained using *n_mute_* = 40, six transformer encoder layers, *d_model_* = 512, and *n_head_* = 8. For models trained during the OpenVaccine competition, *n_head_* is set to 32, *d_model_* is set to 256, dropout is set to 0.1, and conv2d filter size is set to 32, *α* and *β* are both 1. Half of the models were trained with sequences with signal to noise greater than 1 and half were trained with signal to noise greater than 0.5. During post competition experiments, we found that results are better when penalizing measurements with high error more by reducing *α* to 0.5 and increasing *β* to 5. Although using more models at more different conditions can give better results, for better reproducibility we only trained 5 models with *k* = *n_layer_* = 3, 4, 5, 6, 7,8 for each experiment hereinafter. Here *n_head_* is set to 32, *d_model_* is 256, dropout is set to 0.1, and conv2d filter size is set to 8. We also only use sequences with signal to noise greater than 0.25 for training and signal to noise greater than 1 for 10-fold cross validation.

## Author contributions

All authors contributed equally.

## Data availability

Promoter dataset can be downloaded at https://github.com/Shujun-He/Nucleic-Transformer/blob/master/src/promoter_classification/v9d3.csv. The dataset of viral DNA can be downloaded from https://github.com/NeuroCSUT/ViraMiner. OpenVaccine dataset is available at https://www.kaggle.com/c/stanford-covid-vaccine/data. Pretrained models can be accessed from Kaggle notebooks: https://www.kaggle.com/shujun717/nucleic-transformer-promoter-inference, https://www.kaggle.com/shujun717/nucleic-transformer-virus-inference, https://www.kaggle.com/shujun717/nucleic-transformer-rna-degradation-inference.

## Code availability

All training code to fully reproduce results is released at https://github.com/Shujun-He/Nucleic-Transformer, and a web application (Figure S3) developed using H2O.ai’s wave is available at https://github.com/Shujun-He/Nucleic-Transformer-WebApp.

## Acknowledgements

At this time, there is no formal publication detailing the experiments conducted to obtain the OpenVaccine dataset, so we would like to thank Das Lab for providing the data and hosting the OpenVaccine competition.

## Supplementary information

**Figure S1:**
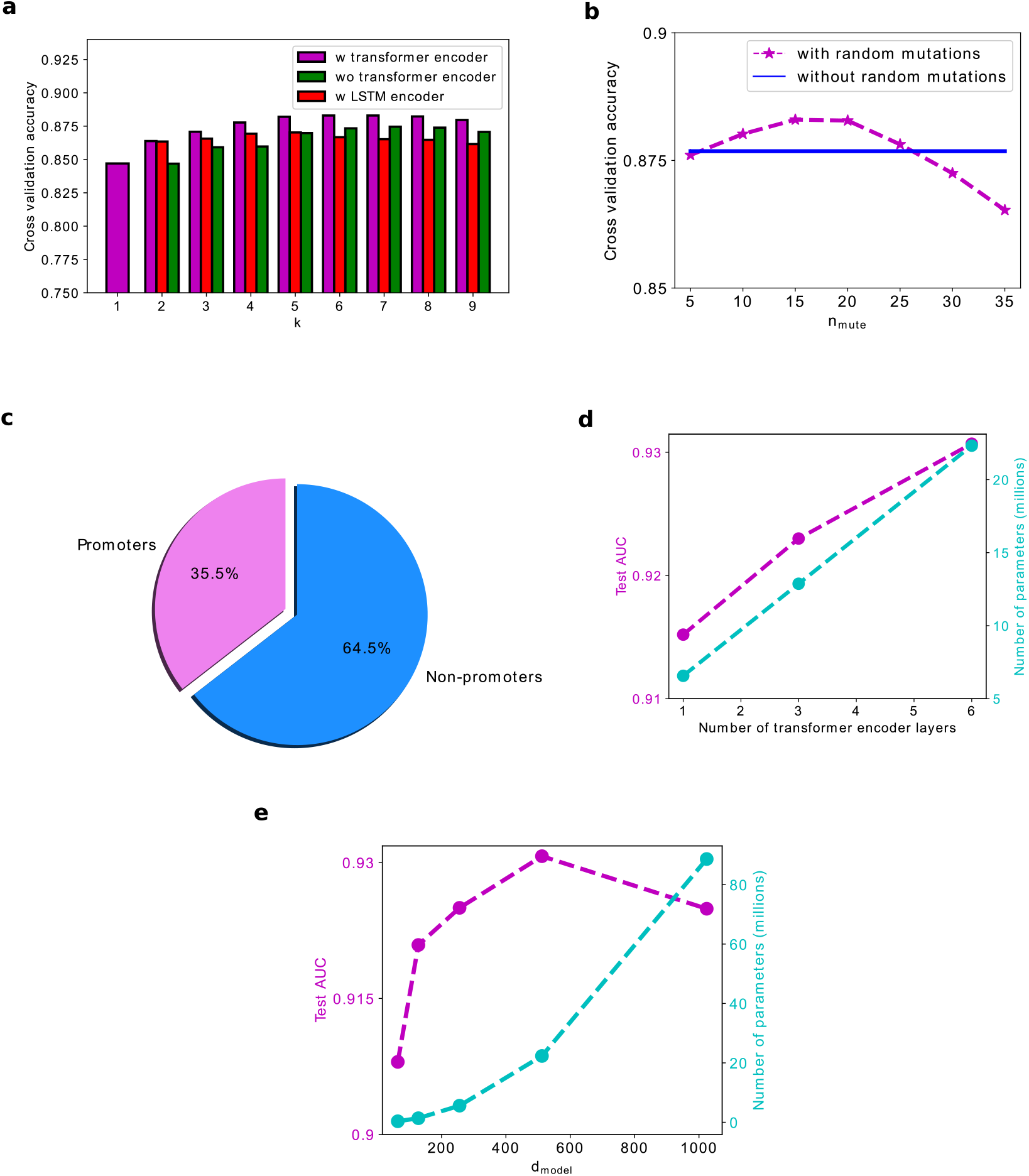
**a.** Bar chart of accuracy vs the parameter *k* of with/without transformer encoder, and with LSTM encoder **b.** Cross validation accuracy vs number of random mutations during training. **c.** Percent of one million randomly generated sequences classified as promoters/non-promoters by the Nucleic Transformer. **d.** The effect of the number of transformer encoder layers on virus test set AUC performance. **e.** The effect of the width of transformer encoder layers on virus test set AUC performance.

**Table S1:** Performance using inputs generated by different biophysical models to predict mRNA degradation in the OpenVaccine dataset.

| Package | Public MCRMSE | Private MCRMSE | Public MCRMSE<br>semi-supervised | Private MCRMSE<br>semi-supervised |
| --- | --- | --- | --- | --- |
| RNAsoft | **0.23154** | 0.34521 | 0.23101 | **0.33841** |
| rnastructure | 0.23637 | 0.34639 | 0.23502 | 0.33989 |
| EternaFold | 0.23158 | 0.34613 | **0.23081** | 0.33878 |
| Contrafold_2 | 0.23245 | 0.34858 | 0.23123 | 0.34111 |
| NUPACK | 0.23679 | 0.34955 | 0.23587 | 0.34292 |
| Vienna | 0.23492 | **0.34515** | 0.2337 | 0.33864 |
| avg of all | 0.22976 | **0.34375** | 0.22914 | **0.33722** |

**Figure S2:**
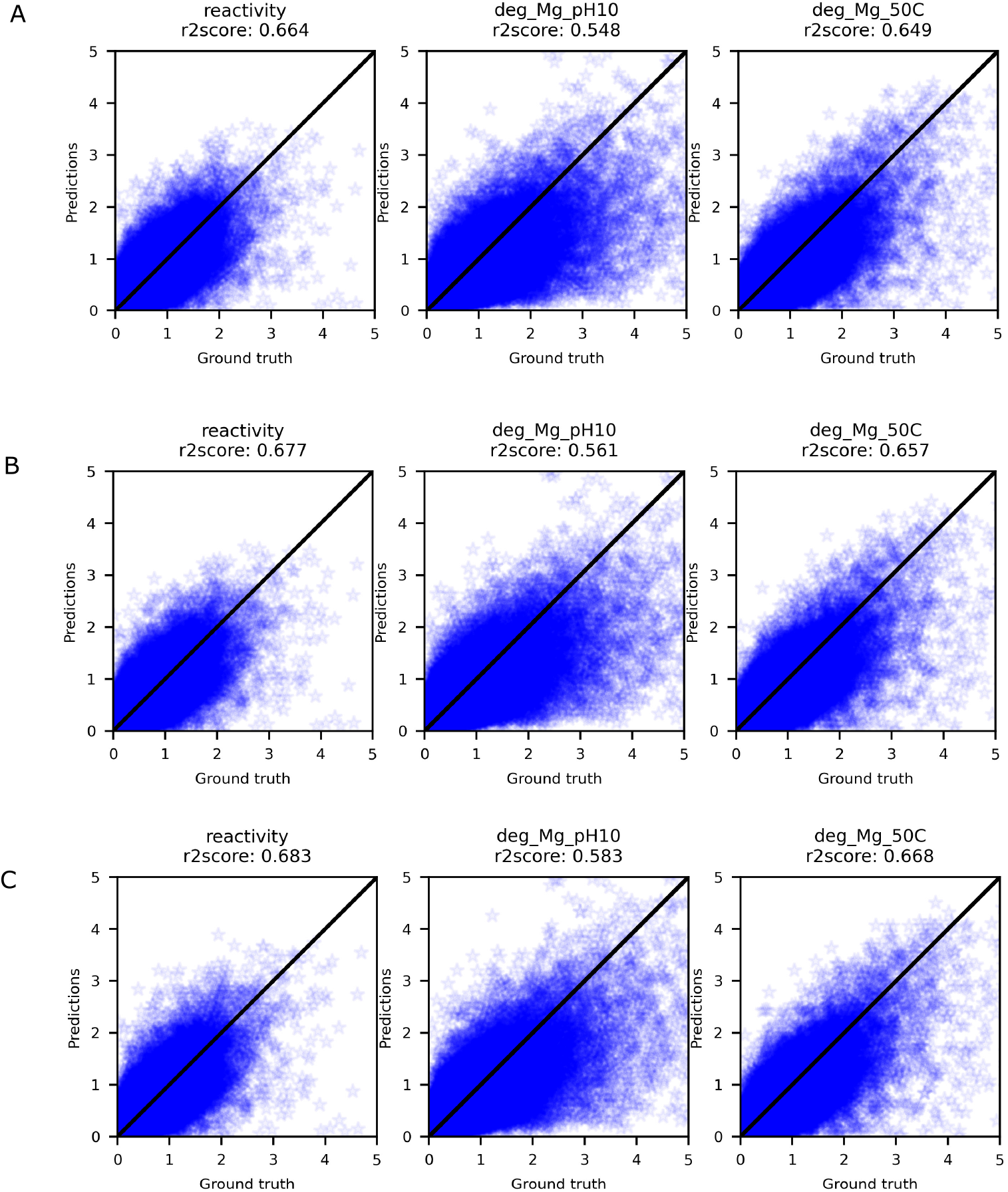
R2 score on the OpenVaccine private test set of the Nucleic Transformer. **a.** supervised only, **b.** unsupervised, **c.** semi-supervised

**Figure S3:**
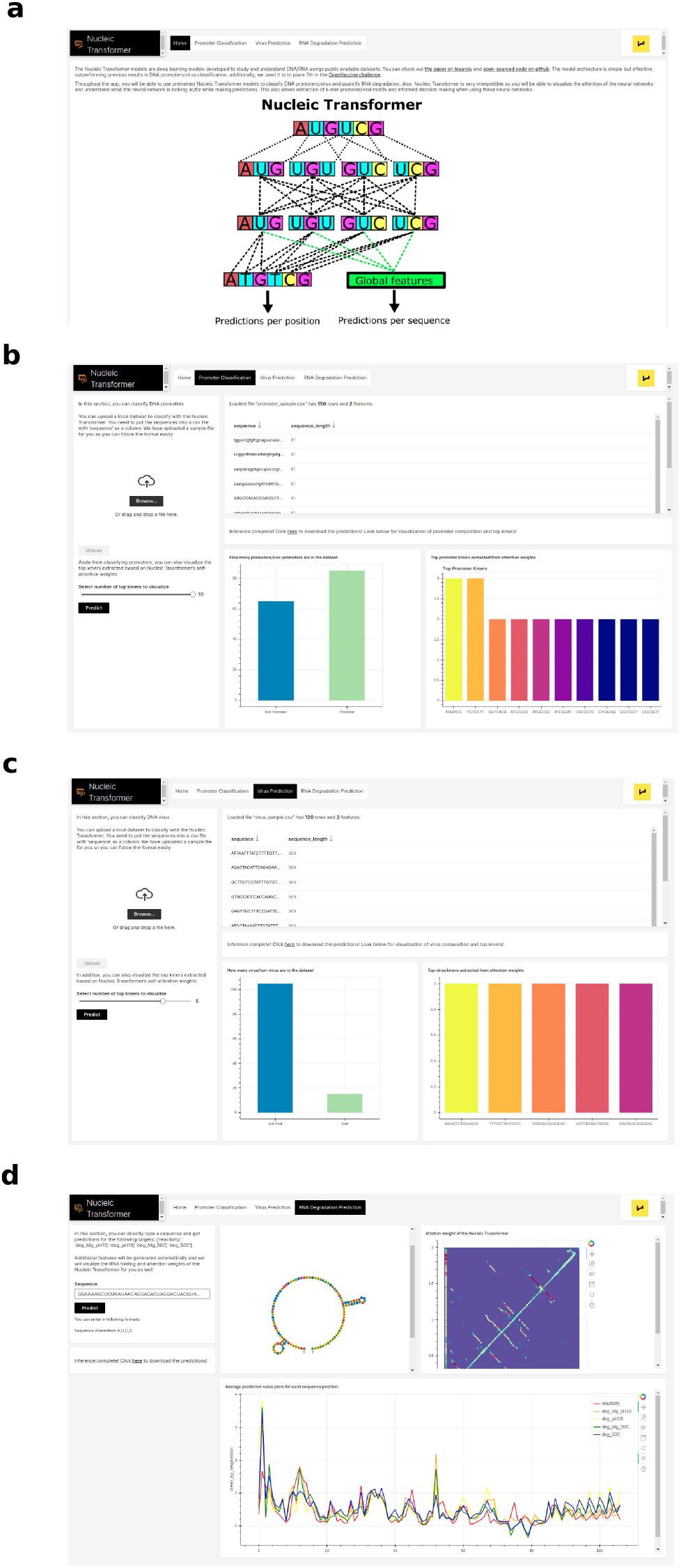
Nucleic Transformer webapp. **a.** Home page **b.** In this page, the user can classify DNA promoters and visualize the top kmers with pretrained models **c.** In this page, the user can classify DNA promoters and visualize the top kmers with pretrained models **d.** In this page, the user can use pretrained models to predict RNA degradation at each nucleotide and visualize the attention weights of the Nucleic Transformer

## Footnotes

1 https://github.com/DasLab/arnie

